# Direct Sputum Sequencing Reveals *Mycobacterium tuberculosis* Diversity Underrepresented by Culture in a High TB-Burden Setting

**DOI:** 10.64898/2026.11.12.766101

**Authors:** B.C. Mann, A. Shaik, A. Gupta, S. Kulkarni, S. Mulaudzi, N. Cesare, K.R. Jacobson, N. Rawoot, R.M. Warren, M.R. Farhat

## Abstract

**Background:** Whole-genome sequencing of *Mycobacterium tuberculosis* (Mtb) commonly relies on culture-derived isolates, potentially altering the bacterial populations represented. Direct sputum whole-genome sequencing (dsWGS) may better capture lineage composition and within-host variant diversity.

**Methods:** We compared targeted-enrichment dsWGS with WGS of matched culture-derived isolates from participants initiating treatment for rifampicin-susceptible pulmonary tuberculosis in Worcester, South Africa. We assessed sequencing performance, technical reproducibility, sputum-sampling variability, lineage and variant concordance, and resistance-associated variation, and explored associations with baseline clinical characteristics and treatment response.

**Results:** Of 218 clinical dsWGS reactions, 139 passed sequencing quality criteria, yielding 113 eligible baseline sputum–culture pairs. Mixed-lineage infection was detected in 72/113 (63.7%) sputum specimens versus 7/113 (6.2%) matched cultures; 66/113 (58.4%) pairs showed reduced lineage diversity following culture. Permutation analysis indicated preferential retention of Lineage 2 and depletion of Lineage 3 after culture. Technical replicates preserved dominant phylotype assignments (median SNP Jaccard similarity 0.863), while independently collected sputum specimens showed greater variability. Among 38 concordant single-lineage pairs, 35,540/38,311 (92.8%) pair-specific variant observations were shared after filtering and raw-read recovery, while 2,497 (6.5%) were sputum-unique and 274 (0.7%) culture-unique. Discordance was concentrated among variable variants; 88.0% of shared variants variable in at least one specimen had higher allele frequencies in culture. No sputum-unique WHO Grade 1 or 2 resistance-associated mutations were identified in the strain-controlled subset, whereas eight mixed-lineage sputum samples versus three matched cultures harboured such mutations. Higher baseline low-frequency variant burden showed non-significant trends towards delayed culture conversion (log-rank p=0.144) and unfavourable treatment outcomes (log-rank p=0.400).

**Conclusions:** Direct-sputum WGS revealed *M. tuberculosis* lineage and low-frequency variant diversity that was not fully represented by matched culture-derived genomes, despite high overall genomic concordance. These findings highlight the complementary value of direct-sputum sequencing for characterising within-host diversity, while suggesting that the representation of bacterial populations following culture may vary across epidemiological settings. Further research is needed to improve the accuracy of minority-variant detection and establish the clinical relevance of diversity measured directly from sputum.

## Introduction

Whole-genome sequencing (WGS) of *Mtb* has become a cornerstone of modern tuberculosis research, enabling high-resolution investigation of transmission, drug resistance, bacterial evolution and within-host *Mtb* diversity (1–4). To date, the vast majority of *Mtb* sequencing have been performed on culture-derived isolates (2, 5, 6). While culture provides a robust and standardized source of high-quality DNA, it introduces biological bottlenecks and selective pressures that may distort the genetic composition of the infecting bacterial population (5–8). Advances in target enrichment technologies have enabled consistent direct from sputum whole-genome sequencing (dsWGS), without the need for culture (2, 5, 9, 10).

Recent studies have demonstrated that dsWGS can identify minority variants or within-sample diversity that may be missed by culture-based sequencing approaches, likely due to culture selection, random loss of low-frequency variants, and adaptation to in vitro conditions (6–8). However, much of this work has focused on technical feasibility, resistance prediction, or selected examples of culture-associated diversity loss, rather than systematic paired comparison of sputum- and culture-derived genomes at scale (2, 9, 11, 12). Moreover, recent work has reported high sputum–culture concordance after careful bioinformatic reconciliation, suggesting that the magnitude of culture-associated diversity loss may be context-dependent (6, 13). Prior datasets provide limited evidence from high-burden, high-transmission settings with substantial HIV co-infection, particularly among patients with drug-susceptible pulmonary TB; therefore, it remains unclear whether culture-associated loss of mixed-lineage and variant diversity occurs at meaningful scale in such settings, or whether these microbiological differences relate to baseline clinical characteristics or treatment response.

In this prospective cohort study, adults initiating treatment for drug-susceptible pulmonary TB in Worcester, South Africa, provided baseline sputum specimens for paired targeted-enrichment dsWGS and culture-derived WGS, with baseline clinical, adherence, culture-conversion, and treatment outcome data collected for exploratory association analyses. Using this paired dataset, we assessed dsWGS feasibility and reproducibility, quantified culture-associated differences in lineage and variant diversity, and explored whether sputum-derived genomic diversity related to clinical or microbiological treatment-response measures.

## Materials and Methods

### Cohort description

Participants initiating treatment for drug-susceptible pulmonary TB were enrolled at the Worcester Community Day Centre (CDC) and Brewelskloof Hospital, South Africa, between 2017 and 2024 (14). Eligible participants were aged 15 years or older and treatment-naive, defined as individuals who had not received therapy for active TB or experienced an active TB episode in the preceding four months. At enrolment, clinical, demographic, radiological, and microbiological data were collected, including age, sex, body mass index (BMI), HIV status, antiretroviral therapy use, previous TB history, incarceration history, alcohol and substance use, household food security, sputum smear microscopy grade, time to positivity (TTP) in liquid culture, and chest radiographic findings, including cavitation and percentage lung involvement.

Sputum sample collection followed a longitudinal design: expectorated sputum specimens were collected once weekly for 12 weeks, with an additional follow-up collection at 5 months. For a subset of participants, a second expectorate (“B sample”) was collected on the same day as the baseline sample to assess within-day variability. All sputum specimens were transported to the Stellenbosch University Biomedical Research Institute for processing, including direct sputum whole-genome sequencing (dsWGS) and culture-based analyses. Participants were followed during treatment and after treatment completion to assess microbiological and clinical outcomes.

Treatment response was assessed using time to sputum culture conversion (TCC) and a composite unfavourable treatment outcome. TCC was defined as the interval from treatment initiation to the first of two consecutive Mtb-negative sputum cultures not followed by a subsequent positive culture. Participants were eligible for TCC analysis if they had at least three uncontaminated sputum specimens and a positive culture within the first five weeks of treatment. The composite unfavourable outcome comprised treatment failure, TB relapse, or death, assessed during treatment and post-treatment follow-up. Deaths attributed to causes unrelated to TB were treated as censored observations. Missing longitudinal culture results were addressed using multiple imputation in the original outcome analysis (14).

### Sample selection, processing and sequencing

Sputum specimens from participants with rifampicin-susceptible pulmonary tuberculosis were selected for direct sputum whole-genome sequencing (dsWGS) using hybrid-capture enrichment. Given the anticipated dependence of sequencing success on bacterial burden, the initial sequencing batch prioritised specimens with higher smear microscopy grades. Subsequent sequencing included specimens across a broader range of smear grades to evaluate performance at lower bacterial burdens, alongside additional specimens selected to assess technical reproducibility and variability between independently collected sputum samples. In total, 218 clinical dsWGS reactions were performed, comprising 174 primary baseline specimens, 12 follow-up specimens, three month-five specimens, 14 additional sequencing reactions from previously extracted DNA to assess technical reproducibility, and 15 independently collected same-day B specimens to assess sampling variability, where A and B denote the first and second expectorated specimens collected at the same study visit, respectively. The technical repeats originated from nine participants. Two *M. tuberculosis* H37Rv reference controls were also included to assess process consistency.

Direct-sputum sequencing was performed using DNA extracted from retained sputum material, while culture-derived WGS was performed on matched isolates obtained from the corresponding clinical specimens. Detailed specimen processing, culture-derived isolate preparation, inactivation, and DNA extraction procedures are described in the Supplementary Methods (Specimen processing and DNA extraction). For paired comparisons, only specimens for which both sequencing datasets met the predefined quality criteria were included. Baseline pairs formed the primary comparison dataset, with follow-up pairs evaluated separately where relevant. Technical repeats and same-day A/B specimens were assessed independently for reproducibility and sampling variability.

Quantitative PCR (qPCR) targeting the single-copy Mtb gene *Rv2341* was used to estimate Mtb genome copy numbers from processed sputum specimens. These estimates informed library pooling during hybrid-capture enrichment and provided an additional measure of bacterial burden for evaluating sequencing performance (10, 15). Assay conditions, primer sequences, and standard-curve quantification are described in the Supplementary Methods (Quantitative PCR). Libraries were prepared using a previously validated custom Twist Bioscience target-enrichment approach (10). Two versions of the capture panel were used, with the modified version reducing probe representation targeting the highly conserved rrs and rrl genes to limit disproportionate read enrichment. Panel-specific sample distributions are provided in Supplementary Data – S1, with library preparation, enrichment and pooling procedures detailed in the Supplementary Methods (Library preparation and target capture). Direct-sputum libraries were sequenced using Illumina MiniSeq or NextSeq 1000/2000 platforms at the Stellenbosch University Centre for Epidemic Response and Innovation (CERI). Culture-derived DNA was sequenced at the Leibniz Lung Research Center, Borstel, Germany.

### Bioinformatics – Core analysis

Sequencing data were processed using a reproducible Snakemake-based bioinformatics workflow (Supplementary Figure - S1). Initial read quality was assessed using FastQC (v0.11.9) (16), followed by adapter trimming and quality filtering with fastp (v0.23.4) (17). Taxonomic classification was performed using Kraken2 (v2.1.3) (18), and reads classified as Mtb were extracted using KrakenTools to reduce interference from non-target sequences (19). Extracted reads were aligned to the H37Rv reference genome (NC_000962.3) using bwa-mem2 (v2.2.1) (20). Secondary and supplementary alignments were excluded, PCR duplicates were removed, and genome coverage and alignment quality metrics were assessed using samtools and Picard Tools (21, 22).

Genome-wide variant calling was performed on deduplicated alignments using FreeBayes (v1.3.6) (23, 24). Variant records underwent decomposition, functional annotation using SnpEff (v5.2) (22, 25), and normalisation using bcftools, with additional corrections to allele-support annotations where required. Variants overlapping predefined low-confidence genomic regions, including repetitive, highly homologous and poorly mappable regions, were excluded using a custom genomic mask. The deduplicated alignments were also analysed independently using TBTypeR (26) for phylogenetic lineage classification. These analyses formed separate downstream branches for genome-wide variant assessment and lineage characterisation. Detailed software parameters, variant-processing procedures and genomic masking criteria are provided in the Supplementary Methods (Bioinformatics - Core analysis).

### Variant quality filtering

Variant quality filtering was performed separately for single-nucleotide polymorphisms (SNPs) and insertions/deletions (indels) to minimise artefactual calls, particularly at low allele frequencies. Candidate SNPs were assessed using read-level alignment and variant-support metrics, including mapping quality, sequencing depth, alternate allele support, strand balance and alignment artefacts. A minimum sequencing depth of 20× was required, with depth-dependent allele-frequency (AF) thresholds applied to account for stochastic sampling effects at lower coverage. Variable SNPs were additionally evaluated using an empirically derived logistic-regression error model, retaining variants with a classification probability ≥0.67, as previously described (14, 23).

Indels underwent separate alignment-based quality filtering, including assessment of read-position biases and adjusted allele frequencies. Filtering criteria were applied as hard thresholds, with variants failing the required criteria excluded from the retained call sets and treated as absent in downstream variant comparisons. This represented an operational classification rather than independent confirmation of the reference genotype. Variants with AF >0.95 were classified as fixed, while those with AF 0.05–0.95 were classified as variable. Detailed SNP- and indel-specific filtering procedures and depth-dependent AF thresholds are provided in the Supplementary Methods (Variant quality filtering; Supplementary Table - S1) (14, 23).

### Lineage classification and concordance analysis

Lineage classification was performed independently of genome-wide variant calling using TBTypeR (v25.01.0) (26). Major lineages and sublineages were identified from curated phylogenetic markers, excluding markers within predefined low-confidence genomic regions. A lineage was considered present when its aggregate allele frequency exceeded 0.05, and mixed-lineage infection was defined as the detection of two or more distinct major lineages within a specimen.

Lineage concordance was evaluated in 113 baseline sputum–culture pairs meeting the predefined sequencing quality criteria. Pairs were assigned to five mutually exclusive categories based on the major lineages detected in each specimen: concordant single-lineage, concordant mixed-lineage, reduced lineage diversity in culture, reduced lineage diversity in sputum, or discordant lineage composition.

To assess whether the lineage composition observed following culture was consistent with lineage-neutral dropout, a permutation-based analysis compared observed culture lineage counts with the distribution expected under random sampling from the direct-sputum lineage pool. The analysis used 200,000 random draws, with the number of sampled lineage observations matched to the total observed in culture. Detailed lineage-calling parameters, concordance definitions and permutation procedures are provided in the Supplementary Methods (Lineage classification and concordance analysis).

### Variant concordance analysis

Variant concordance was assessed in 38 baseline sputum–culture pairs with concordant single-major-lineage classifications, where both sequencing datasets met the predefined quality criteria. Technical replicates were excluded. Within each pair, quality-filtered variants were matched by genomic position, reference allele and alternate allele, and initially classified as shared, direct sputum-unique or culture-unique. A bidirectional recovery step examined pre-filter sequencing evidence in the matched specimen at positions of initially unique variants. Variants with qualifying alternate-allele evidence were recovered and reclassified as shared, while those remaining detectable in only one specimen were retained as specimen-unique.

Concordance was summarised using pair-specific variant observations, with shared variants considered fixed only when fixed in both specimens. Total variant counts were compared between specimen types, and allele-frequency shifts among shared variants were calculated as ΔAF = AF_direct sputum − AF_culture. Paired comparisons were assessed using Wilcoxon signed-rank tests, with allele-frequency analyses evaluated at the specimen-pair level. Detailed recovery criteria and concordance procedures are provided in the Supplementary Methods (Variant concordance and recovery).

### Genomic distribution and annotation-level analysis of specimen-unique variants

The genomic distribution of specimen-unique variants was examined across the 38 concordant single-lineage baseline sputum–culture pairs following variant filtering and bidirectional raw-read recovery. Direct sputum-unique and culture-unique variant observations were mapped to their genomic positions and corresponding gene or genomic-region annotations. Genome-wide distributions and the most frequently represented annotations were examined separately for each specimen type to identify regions contributing to differences in detectable variation. Detailed analytical procedures are provided in the Supplementary Methods (Genomic distribution and annotation-level analysis of specimen-unique variants).

### Drug-resistance-associated variant analysis

Drug-resistance-associated variation was investigated separately in concordant single-lineage and mixed-lineage infections. In the 38 strain-controlled baseline sputum–culture pairs, variants within a curated set of drug-resistance-associated genes were assessed for concordance following variant filtering and bidirectional raw-read recovery. Distinct mutations were evaluated against the WHO drug-resistance mutation catalogue using exact genomic position, reference allele and alternate allele matching (27). An exploratory analysis was additionally conducted in 72 baseline sputum–culture pairs classified as mixed-lineage by direct sputum sequencing. Quality-filtered variants from each specimen type were screened independently for WHO Grade 1 and Grade 2 resistance-associated mutations, and their detection and allele frequencies were compared between matched specimens. Detailed procedures for both analyses are provided in the Supplementary Methods (Drug resistance associated variant analysis).

### Technical reproducibility, sputum sampling variability, and process controls

Technical reproducibility was assessed using independently prepared and sequenced libraries generated from the same sputum DNA extract. Seven specimen sets met the sequencing quality criteria for replicate analysis, comprising 18 sequencing runs and 15 pairwise comparisons. Reproducibility was evaluated at the variant and phylogenetic levels by comparing quality-filtered SNP and indel profiles, major-lineage assignments and TBTypeR phylotype classifications. Variant-level agreement was quantified using Jaccard similarity, with separate assessment of fixed and variable variants.

To evaluate variability between independently collected specimens, same-day A/B sputum pairs were compared using major-lineage and TBTypeR phylotype classifications. Six pairs in which both specimens met sequencing quality criteria were included. Two H37Rv reference process controls were additionally evaluated for consistency in genomic variant profiles and evidence of unexpected lineage signals. Detailed replicate-comparison procedures and similarity calculations are provided in the Supplementary Methods (Technical reproducibility, sputum sampling variability, and process controls).

Longitudinal variant concordance was assessed in three participants with baseline and follow-up direct-sputum specimens that both met the predefined sequencing quality criteria. Fixed SNPs, error-model-filtered variable SNPs (AF 0.05–0.95), and quality-filtered fixed and variable indels were compared between time points. Concordance was summarised using the number of shared and time-point-specific variants and Jaccard similarity.

### Exploratory clinical correlates of mixed-lineage infection

Associations between mixed-lineage infection and baseline clinical characteristics were investigated in 113 participants with confirmed paired baseline direct-sputum and culture WGS results. Participants were classified according to their baseline direct-sputum lineage composition as mixed-lineage (≥2 major lineages) or single-lineage (one major lineage). Clinical, demographic, behavioural, microbiological and radiological variables were obtained from the updated TRUST II clinical dataset and linked at the participant level. Categorical variables were compared using Pearson’s chi-square tests without continuity correction, and continuous variables using two-sided Mann–Whitney U tests. Analyses were performed using available observations for each variable, with baseline liquid-culture time to positivity assessed using the recorded whole-day value for the first sputum specimen. All comparisons were exploratory, and p-values were reported without adjustment for multiple testing. Detailed variable definitions and results are provided in the Supplementary Methods and Supplementary Table – S5.

### Low-frequency variant burden and treatment-response analysis

The association between baseline low-frequency (LF) variant burden and treatment response was assessed in the 38 concordant single-lineage baseline participants described above. LF variants were defined as variants with an allele frequency of 0.05–0.95, with the total LF burden calculated as the sum of error-model-filtered SNPs and quality-filtered indels detected in each direct sputum specimen. Treatment response was assessed using the previously derived TRUST II time-to-culture-conversion (TCC) and composite unfavourable treatment-outcome variables, as defined above.

Participants were grouped by baseline LF variant burden (>25 versus ≤25 variants), a threshold that minimised the AIC across candidate thresholds. LF burden was also analysed as a continuous log₂-transformed variable, and a median split (>31 variants) was used as an alternative threshold. Kaplan-Meier curves and log-rank tests compared outcomes between groups, and restricted mean survival time to 12 weeks was estimated for TCC. Univariable Cox proportional hazards models estimated hazard ratios and 95% confidence intervals.

An exploratory multivariable Cox model was fitted for TCC, adjusted for percent lung involvement, previous tuberculosis, time to positivity (per day), age, body mass index <18 kg/m², smoking and diabetes, and stratified by HIV status, lung involvement (≤25% vs >25%) and sex, with ridge penalisation (λ=0.1) to reduce model instability. Smear status was not included because 29 of 30 participants with TCC data were smear-positive. Sensitivity analyses excluded S0224_01 (approximately 50/50 lineage 2 mixture) and S0186_01 (low-input outlier). Analyses were restricted to participants with available outcome data.

### Accession number(s)

Sequencing data generated in this study will be deposited in the NCBI Sequence Read Archive, with the BioProject accession number provided upon publication.

### Ethics

Ethical approval for this study was obtained from the South African Medical Research Council Ethics Committee under reference number EC011-5/2016 and subsequently also from the Stellenbosch University Health Research Ethics Committee (HREC) under reference number N17/04/039 RECIP_MRC_EC011-5/2016. The protocol was also approved by the Boston Medical Center and Harvard Medical School Institutional review boards that deferred to the Stellenbosch IRB for full review.

## Results

### Cohort characteristics and direct performance

Between 2017 and 2024, we enrolled 458 participants initiating treatment for rifampicin susceptible pulmonary tuberculosis (Table 1). The cohort was predominantly male (60.5%), with a median age of 37 years (IQR 26–48), and a high prevalence of HIV co-infection (28%) and prior TB history (41%). The median time to positivity (TTP) in liquid culture was 6 days, and 48% of participants were smear grade 1+ or higher, indicating a high average bacillary burden (Table 1).

**Table 1.** Baseline demographics of the full TRUST cohort, paired baseline dsWGS–culture subset, and concordant single-lineage paired subset.

| Variable | Full TRUST cohort (N=458) | Paired baseline dsWGS–culture subset (N=113) | Concordant single-lineage paired subset (N=38) |
|---|---|---|---|
| Sex |  |  |  |
| Male | 277 (60.5%) | 66 (58.4%) | 24 (63.2%) |
| Female | 181 (39.5%) | 47 (41.6%) | 14 (36.8%) |
| Age, years, median (Q1–Q3) | 37 (26–48) | 37 (25–47) | 37.5 (25.5–45.8) |
| BMI, kg/m², median (Q1–Q3) | 18 (16–20) | 17.6 (15.8–19.6) | 17.6 (15.4–19.8) |
| Smear grade |  |  |  |
| No AFB | 120 (33.8%) | 14 (12.4%) | 3 (7.9%) |
| Scanty | 65 (18.3%) | 7 (6.2%) | 1 (2.6%) |
| 1+ | 42 (11.8%) | 26 (23.0%) | 8 (21.1%) |
| 2+ | 52 (14.6%) | 20 (17.7%) | 10 (26.3%) |
| 3+ | 76 (21.4%) | 46 (40.7%) | 16 (42.1%) |
| HIV positive | 128 (28.0%) | 28 (24.8%) | 6 (15.8%) |
| Prior TB | 187 (40.8%) | 49 (43.4%) | 15 (39.5%) |
| TTP, days, median (Q1–Q3) | 6 (5–10) | 6 (4–8) | 5 (4–6) |
| PLI, %, median (Q1–Q3) | 21.5 (13.8–30.3) | 23.8 (15.1–32.4) | 28.5 (20.5–37.4) |

Data are presented for the full TRUST cohort, the baseline paired direct sputum WGS and culture-derived WGS analysis set, and the subset of baseline pairs in which both direct sputum and culture were classified as single-lineage and assigned to the same major lineage. The concordant single-lineage paired subset was used for strain-controlled variant-level comparisons. Values are n (%) or median (Q1–Q3).

To capture the population characteristics of these infections prior to in vitro selection, we performed targeted hybrid-capture direct-from-sputum whole-genome sequencing (dsWGS). We sequenced 218 clinical sputum-derived dsWGS reactions alongside two *M. tuberculosis* H37Rv process controls. Among the clinical dsWGS reactions, 139 met our predefined quality threshold of ≥95% genome coverage at ≥20× depth. After pairing QC-passing direct-sputum results with eligible same-specimen culture-derived WGS data, 116 matched sputum–culture comparisons were available for downstream analyses, comprising 113 baseline pairs and three follow-up pairs. Same-day B samples, technical repeat sequencing runs, month-5 samples, and cases where only a later culture time point was available were retained for separate analyses where relevant but were not counted as paired baseline sputum–culture comparisons. Specifically, six same-day A/B sample pairs had both specimens pass QC, seven baseline samples had an evaluable technical repeat set with the original and at least one repeat passing QC, and none of the three Month-5 samples met the dsWGS QC threshold.

In multivariable logistic regression using non-repeat clinical dsWGS samples with positive quantitative bacterial-load estimates, both smear grade and quantitative bacterial load independently predicted sequencing success. Each ordinal increase in smear grade approximately doubled the odds of dsWGS success (OR 2.09, 95% CI 1.62–2.69; p < 0.001), while each log₁₀ increase in Mtb genome copies increased the odds of success (OR 1.45, 95% CI 1.05–2.01; p = 0.025). Sequencing failures were concentrated among lower-burden specimens: 55/79 failures were smear-negative or scanty, and 61/79 yielded fewer than 10 million Mtb-classified reads. Among QC-passing clinical dsWGS reactions, the mean sequencing depth was 246.1× ± 289.5×. The proportion of reads classified as Mtb by Kraken2 after capture was 82.6% ± 18.3%, while host-derived reads accounted for 3.6% ± 3.7% on average (Supplementary Figure – S2).

### Direct sputum sequencing reveals substantial mixed-lineage diversity purged by culture

We next compared the phylogenetic composition of 113 eligible paired baseline sputum and culture genomes. dsWGS revealed a strikingly higher prevalence of mixed-lineage infections: two or more major lineages were detected in 63.7% (72/113) of direct sputum samples compared to only 6.2% (7/113) of matched culture-derived genomes. In sputum-culture sequence concordance assessment, reduced lineage diversity in culture was the dominant pattern, observed in 58.4% (66/113) of pairs. By contrast, 6/113 (5.3%) were concordant mixed-lineage, 3/113 (2.7%) were discordant and 33.6% (38/113 pairs) of pairs were concordantly single-lineage, highlighting that standard *in vitro* culture routinely fails to capture the true polyclonal complexity of *in vivo* infections in our high TB prevalence setting (Figure 2).

**Figure 2:**
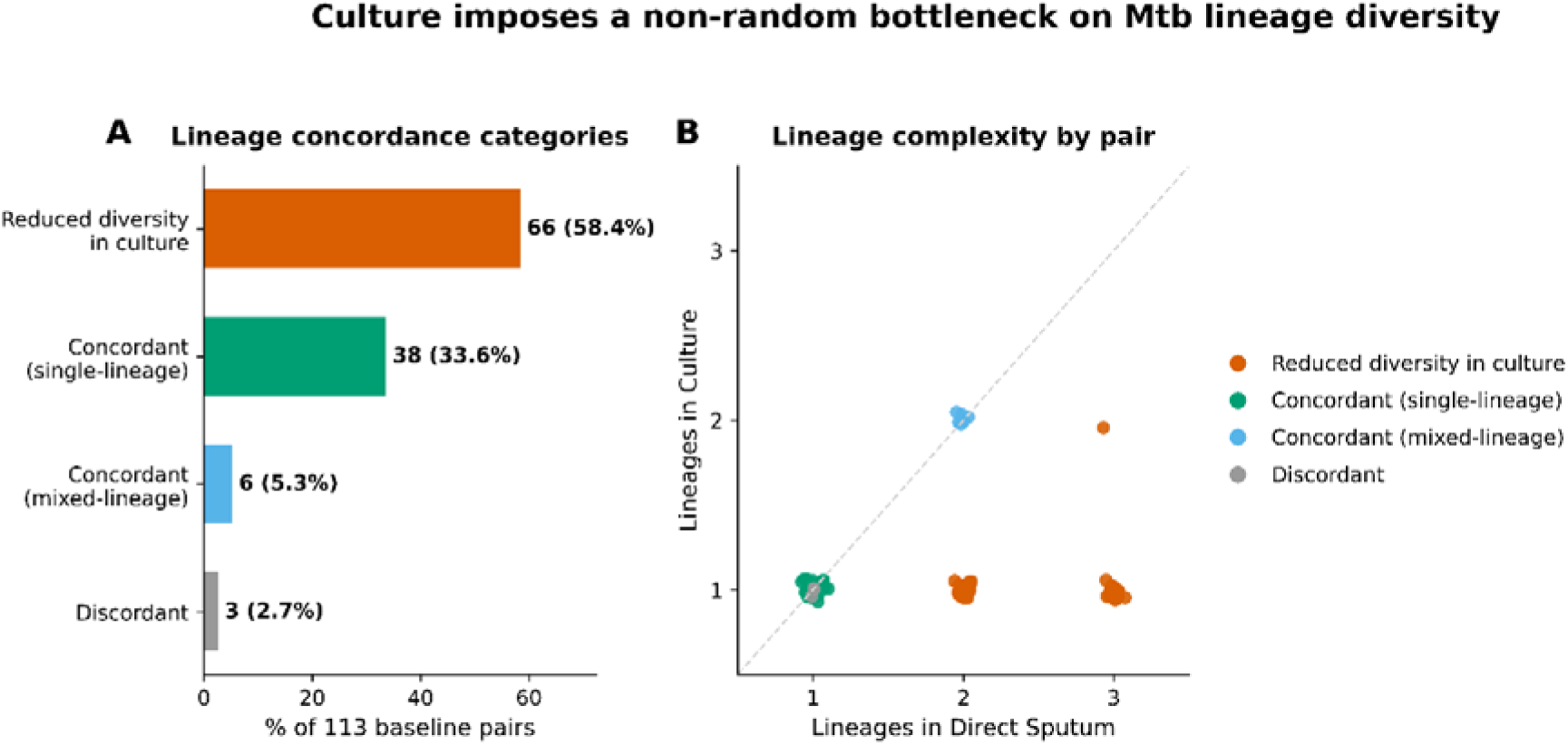
Culture-associated reduction in Mtb lineage diversity. (A) Distribution of paired lineage-concordance categories among 113 eligible baseline direct sputum– culture pairs. Reduced diversity in culture was the dominant pattern, followed by concordant single-lineage pairs, concordant mixed-lineage pairs, and discordant pairs. No baseline pairs showed reduced diversity in sputum. (B) Paired comparison of lineage complexity, showing the number of major lineages detected in direct sputum versus the corresponding culture-derived genome.

To assess whether the lineage composition recovered after culture was consistent with lineage-neutral dropout, we compared observed culture lineage counts with the counts expected under random sampling from the direct-sputum lineage pool. Across the 113 baseline pairs, direct sputum contained 208 major-lineage observations compared with 120 culture observations. In a permutation test using 200,000 draws, L2 was recovered in culture more often than expected under neutral sampling (65 observed vs 54.8 expected; permutation p ≈ 0.005), whereas L3 was depleted (6 observed vs 15.0 expected; permutation p < 0.001). L4 did not differ from neutral expectation (49 observed vs 50.2 expected; permutation p ≈ 0.78). These findings suggest that culture-associated lineage loss was not random, with preferential retention of L2 and depletion of L3, although lineage-specific estimates should be interpreted cautiously because some lineages, particularly L3, were represented by fewer observations.

### Direct sputum sequencing is technically reproducible but sampling dependent

H37Rv process controls showed broadly consistent variant profiles, with no coherent secondary-lineage signals suggestive of substantial cross-contamination. Technical replicate sequencing across seven participants (18 sequencing runs) demonstrated consistent recovery of dominant phylotypes across all 15 pairwise comparisons, with a median SNP Jaccard similarity of 0.863. Agreement was lower for variable SNPs (median Jaccard = 0.581), indicating greater variability in individual lower-frequency variant detection despite reproducible dominant genomic assignments (Supplementary Table – S2).

Independent same-day A/B sputum specimens showed variation in lineage composition. Among six pairs passing dsWGS quality control, four had identical major-lineage profiles (mean Jaccard = 0.833), while three had identical complete TBTypeR phylotype sets (mean Jaccard = 0.750, allowing hierarchical matching of related phylotype labels). Differences included shifts in relative lineage proportions and changes in minority phylotype detection. Together, these findings support reproducible recovery of dominant genomic populations from the same DNA extract while highlighting additional variability associated with independent sputum sampling (Supplementary Table - S3).

Longitudinal variant concordance was additionally assessed in three participants with QC-passing baseline and follow-up direct-sputum specimens. Fixed SNP profiles were relatively conserved between time points (Jaccard similarity 0.877–0.994), with 651–1,041 shared fixed SNPs per pair. In contrast, concordance among error-model-filtered low-frequency SNPs was substantially lower (Jaccard similarity 0.023–0.111), with only 2–5 shared variants per pair. Filtered indels showed intermediate concordance (Jaccard similarity 0.261–0.450). These findings suggest relative stability of the dominant genomic variant profile alongside greater variability in detectable low-frequency variation, although interpretation is limited by the small number of longitudinal pairs and potential differences in sequencing depth and sputum sampling.

### Direct sputum sequencing reveals additional low-frequency variation despite high genome-wide concordance with culture

To assess differences in variant repertoires independently of major-lineage composition, we restricted our analysis to 38 concordant single-lineage baseline sputum–culture pairs. Following variant filtering and bidirectional raw-read recovery, 38,311 pair-specific variant calls were classified, of which 35,540 (92.8%) were shared, 2,497 (6.5%) were unique to direct sputum, and 274 (0.7%) were unique to culture. Median variant counts were 1,147 in direct sputum and 1,129 in culture (paired Wilcoxon p < 0.001), with variation in the extent of discordance between individual pairs (Figure 3).

**Figure 3:**
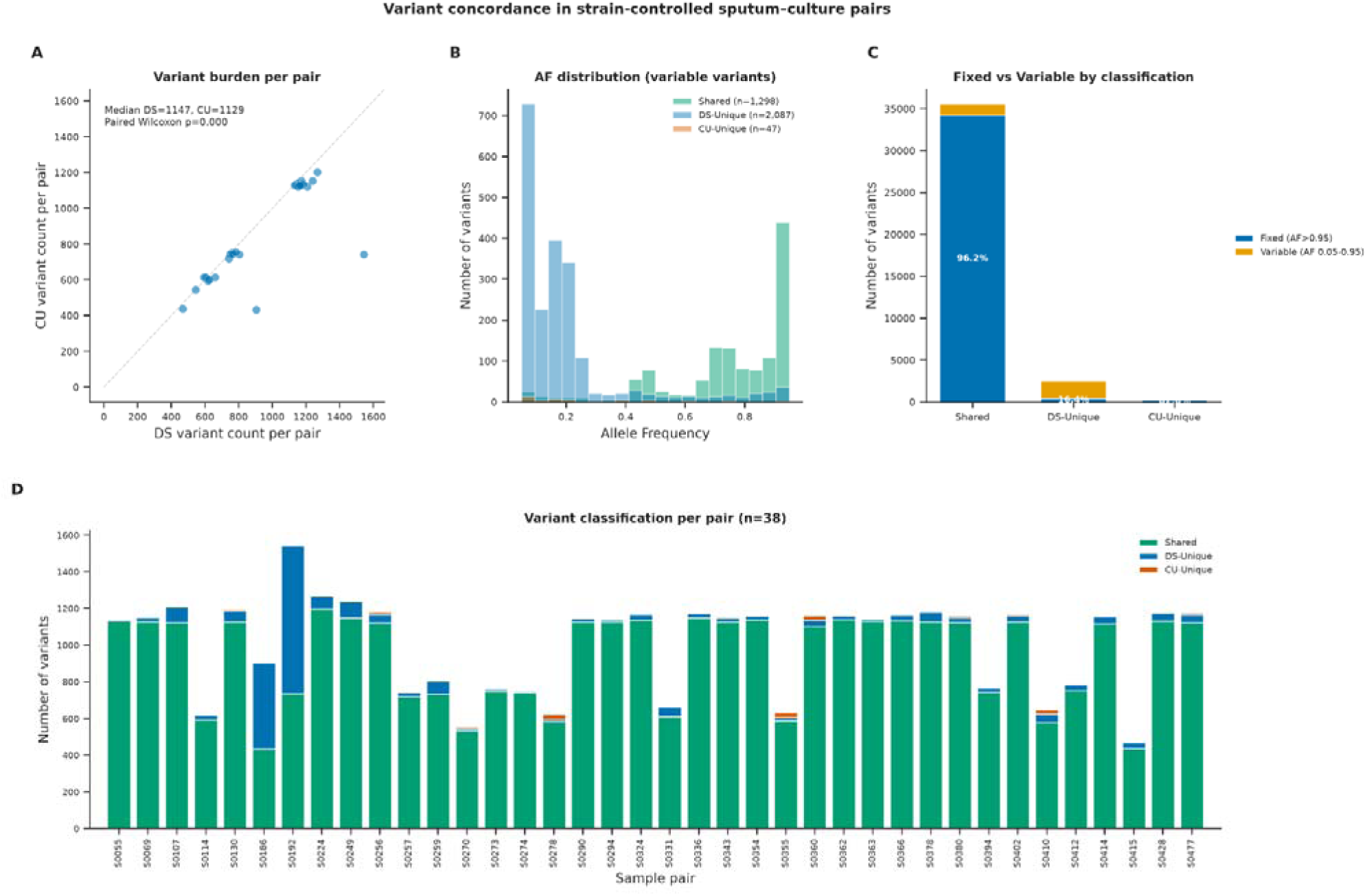
Genome-wide variant concordance between direct sputum and culture-derived WGS in 38 strain-controlled baseline pairs. (A) Paired comparison of total variant counts. (B) Allele-frequency distributions of variable variants stratified by concordance classification. (C) Proportions of fixed and variable variants within each classification; shared variants were considered fixed only when fixed in both specimens. (D) Distribution of shared, direct sputum-unique and culture-unique variants across individual specimen pairs.

Differences in variant recovery were concentrated among variable variants (AF 0.05– 0.95). Of the sputum-unique calls, 2,087/2,497 (83.6%) were variable, compared with 47/274 (17.2%) culture-unique calls. In contrast, 34,204/35,540 (96.2%) shared variants were fixed in both specimens. Among shared variants, 23.9% had higher allele frequencies in culture, 6.1% in direct sputum, and 70.0% were unchanged (Supplementary Figure – S3). This difference was more pronounced among the 1,336 shared variants variable in at least one specimen, of which 88.0% had higher allele frequencies in culture (median ΔAF [direct sputum − culture] = −0.112; pair-level Wilcoxon p < 0.001). Together, these findings demonstrate high overall genome-wide concordance while highlighting additional low-frequency variation and differences in allele-frequency structure between specimen types.

### Genomic distribution and gene-level patterns of specimen-unique variants

Exploratory gene-level analyses were performed to characterise variants uniquely detected in direct sputum or culture-derived WGS. The 2,497 sputum-unique and 274 culture-unique variant observations were distributed across the reference genome, although several genomic regions contributed disproportionately to the observed discordance (Supplementary Figure – S4). The most frequently represented sputum-unique annotations included *fadE23-fadB4* (75 observations), *icl1-fadB2* and *Rv3647c-cspA* (57 each), whereas culture-unique observations were most frequently associated with *prfB-fprA* (24) and *Rv1668c* (23).

### Direct sputum sequencing captures drug-resistance-associated variants not recovered in culture

To determine whether specimen-specific variant differences affected genomic regions relevant to drug resistance, we examined variants in drug-resistance-associated genes among the 38 concordant single-lineage baseline sputum–culture pairs. Across 13 genes containing observed variants, 411 pair-specific variant observations were identified, comprising 395 (96.1%) shared between direct sputum and culture, 16 (3.9%) detected only in direct sputum, and none detected only in culture (Figure 4).

**Figure 4:**
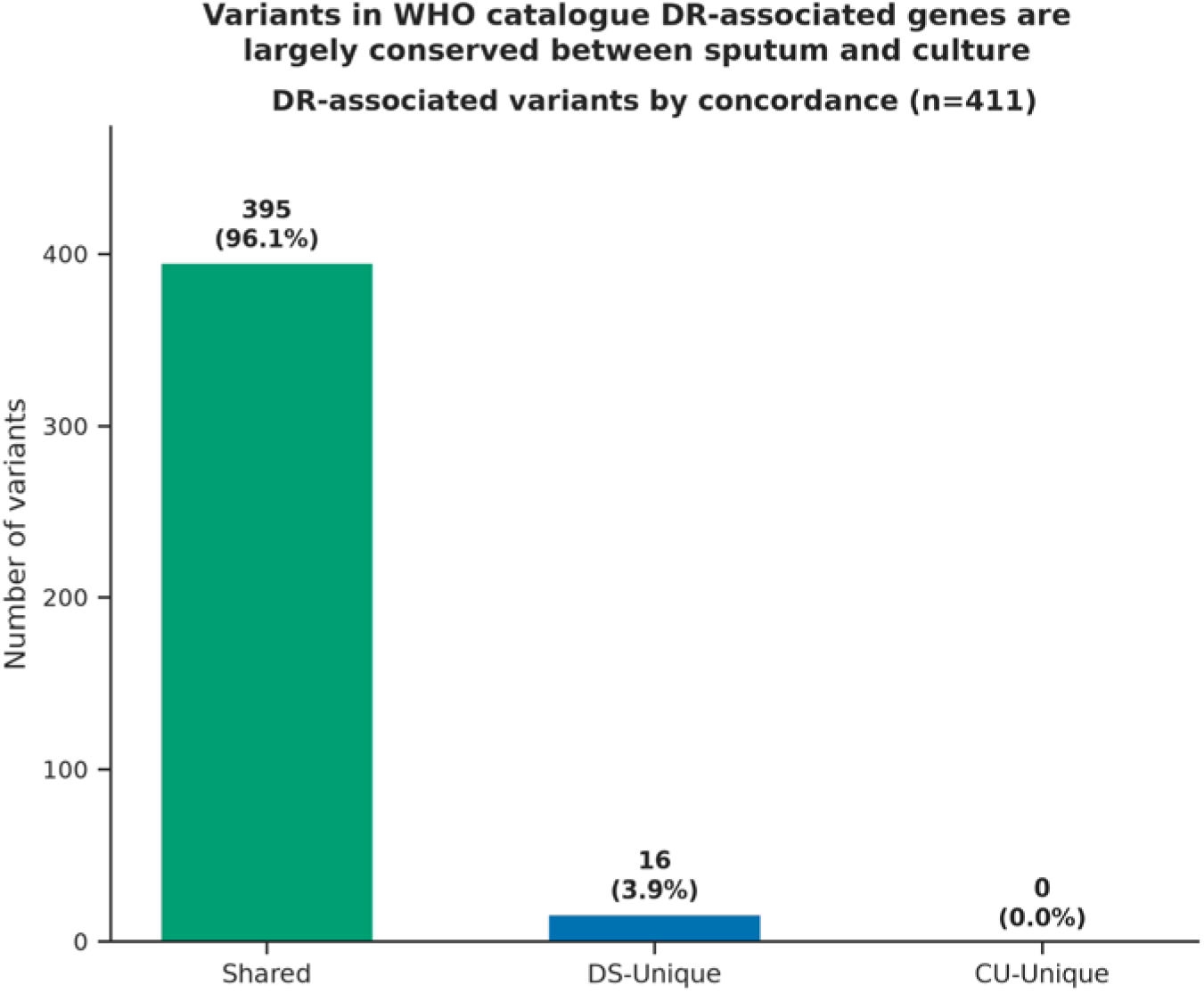
Concordance of variants in drug-resistance-associated genes between direct sputum and culture-derived whole-genome sequencing. Classification of 411 pair-specific variant observations identified across 38 concordant single-lineage baseline sputum–culture pairs. Variants were classified as shared, direct sputum-unique or culture-unique. Variant counts represent observations across paired specimens rather than distinct mutation identities.

The 16 sputum-unique observations represented eight distinct genomic variants across seven specimen pairs, with a median sputum allele frequency of 0.215. These observations occurred in *Rv1258c* (n = 5), *gid* (n = 3), *Rv0194*, *katG* and *embB* (n = 2 each), and *gyrA* and *eis* (n = 1 each). Exact mutation-level comparison with the WHO 2026 drug-resistance mutation catalogue identified no Grade 1 or Grade 2 resistance-associated variants. One recurrent *Rv1258c* frameshift was classified as having uncertain significance for pyrazinamide resistance (Grade 3), while other classified sputum-unique variants were designated as not associated with resistance (Grade 5). Together, these findings indicate high sputum–culture concordance within the examined drug-resistance-associated regions, without evidence of confidently resistance-associated mutations uniquely detected in sputum.

Among 72 baseline sputum–culture pairs classified as mixed-lineage by direct-sputum sequencing, WHO Grade 1 or 2 resistance-associated variants were detected in eight sputum specimens (11.1%) and three matched culture-derived specimens (4.2%). Overall, 19 specimen-specific genomic variant records representing nine distinct genomic variants were identified in direct sputum. Of these, five (26.3%) were also detected in matched culture, whereas 14 (73.7%) were detected only in sputum. No WHO Grade 1 or 2 variants were detected exclusively in culture. All 14 sputum-only variant records occurred in pairs exhibiting reduced lineage diversity following culture, whereas four of the five shared records occurred in pairs retaining mixed-lineage status (Supplementary Table - S4). These findings are consistent with reduced detection of resistance-associated variation following culture, although differences in variant recovery cannot be attributed exclusively to changes in strain composition.

### Exploratory clinical correlates of mixed-lineage infection

Baseline clinical characteristics were compared between 72 participants with mixed-lineage infection (63.7%) and 41 with single-lineage infection (36.3%). Problem alcohol use was more prevalent among participants with mixed-lineage infection (66.7% versus 39.0%; p = 0.004). The distribution of baseline phosphatidylethanol (PEth) categories also differed between groups (p = 0.020), with fewer mixed-lineage participants having PEth concentrations below 50 ng/mL (47.2% versus 71.8%). Mixed-lineage infection was associated with longer liquid-culture time to positivity (median 6 days, IQR 5–8, versus 5 days, IQR 4–6; p = 0.034) and lower radiographic percentage lung involvement (median 20.8%, IQR 13.9–28.8, versus 27.2%, IQR 19.1–37.7; p = 0.009). No clear differences were observed in other evaluated characteristics, including sex, HIV status, previous tuberculosis, body mass index category or baseline smear positivity. These unadjusted associations were exploratory and should be interpreted as hypothesis-generating (Supplementary Table – S5).

### Low-frequency variant burden and treatment response in single-lineage infections

LF variant counts were significantly higher in mixed-lineage than in single-lineage infections (median 910, IQR 625–1038.5, among 72 mixed-lineage patients vs median 24, IQR 16.5–44.25, among 40 single-lineage patients with an available LF count; Mann–Whitney p=7.2×10⁻¹). This likely reflects the contribution of variants from co-circulating strains, which cannot readily be distinguished from within-strain LF variants in mixed infections. All subsequent treatment-response analyses were therefore restricted to the 38 baseline patients with concordant single-lineage infections.

Among these 38 patients, all had at least one LF variant (range 1–1,130), with a median of 24 variants (IQR 15.5–42.25). The primary lineage distribution was lineage 2 (n=23, 61%), lineage 4 (n=14, 37%) and lineage 3 (n=1, 3%). LF variant burden was not significantly associated with smear grade among smear-positive patients with a recorded grade (Spearman r=0.03, p=0.853, n=34), percent lung involvement (r=0.00, p=0.995), TTP (r=0.14, p=0.410), or infecting lineage (lineage 2 median 25 variants, IQR 18.5–45.5 vs lineage 4 median 20 variants, IQR 14.25–38.75; Mann-Whitney p=0.661). A modest inverse association was observed with quantitative bacterial load (Spearman r=−0.40, p=0.012 between log₁₀(Mtb genome copies) and log₁₀(LF variant count)), indicating that patients with higher bacterial burden at baseline tended to have fewer detectable LF variants.

Of the 38 patients, 30 had TCC data (18 events) and 34 had composite unfavourable outcome data (7 events). Participants were grouped by baseline LF variant burden (>25 versus ≤25 variants), a threshold that minimised the AIC across candidate thresholds. Of the 38, 18 had more than 25 variants and 20 had 25 or fewer (16 and 14 among those with TCC data, 18 and 16 among those with composite outcome data). In a Kaplan-Meier analysis, LF variant burden above 25 was not significantly associated with culture conversion (log-rank p=0.144; Figure 5A); 8 of 16 high-burden patients converted compared with 10 of 14 low-burden patients, and the restricted mean time to conversion over 12 weeks was 1.06 weeks longer in the high-burden group. In univariable Cox analysis, the HR for culture conversion was 0.52 (95% CI 0.20–1.32, p=0.171), indicating a tendency towards slower conversion at higher burden. In the exploratory multivariable model, the HR was 0.33 (95% CI 0.10–1.11, p=0.072; n=30, 18 events), and no covariate reached statistical significance. LF burden analysed as a continuous log-transformed variable was not associated with culture conversion (HR 0.98, p=0.819). LF variant burden above 25 was not significantly associated with the composite unfavourable outcome (log-rank p=0.400): 6 of 18 high-burden patients had an unfavourable outcome compared with 1 of 16 low-burden patients (HR 2.55, 95% CI 0.27–24.56, p=0.417; 7 events among 34 patients). The estimate was imprecise given the small number of events, and the continuous log analysis showed no association (HR 1.03, p=0.905).

**Figure 5.**
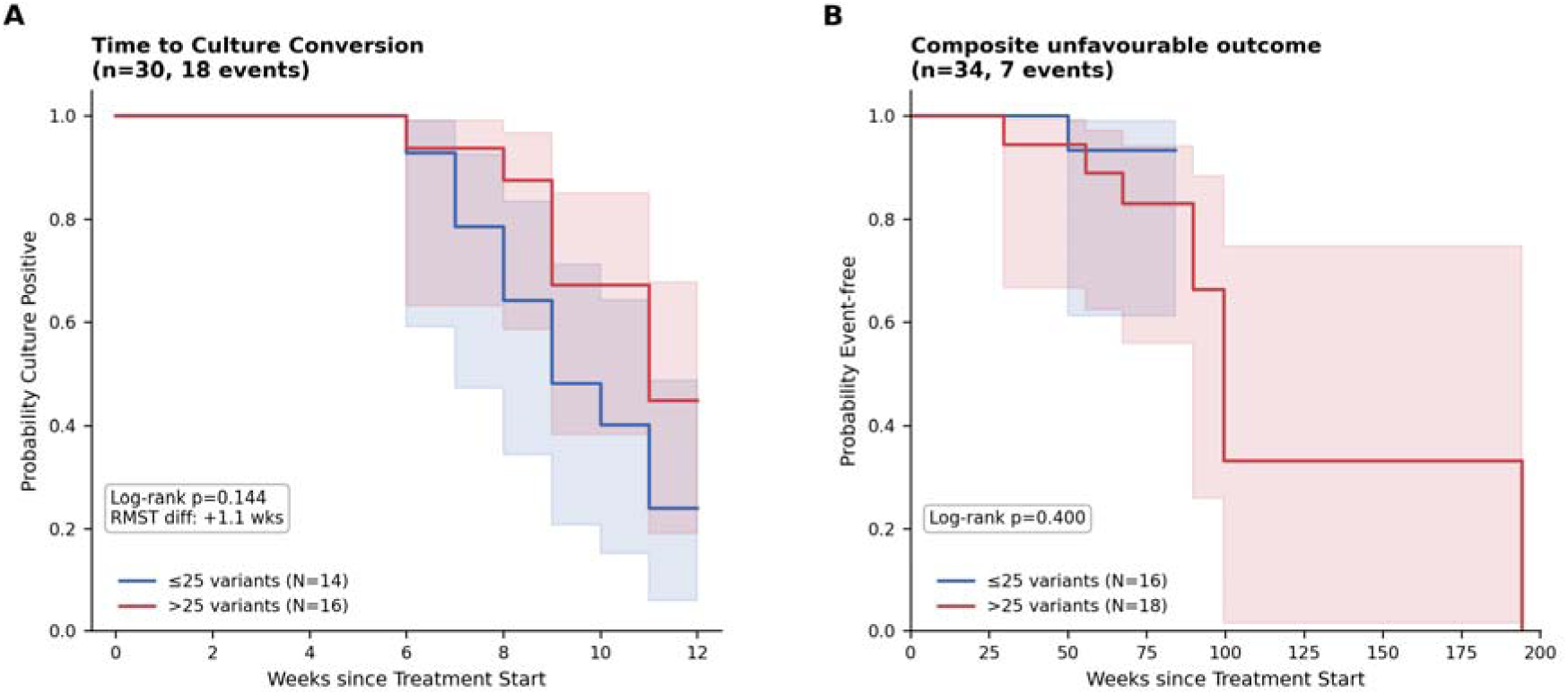
Baseline low-frequency variant burden and treatment response in concordant single-lineage infections. Kaplan-Meier curves comparing participants with more than 25 versus 25 or fewer low-frequency (LF) variants at baseline. (A) Time to culture conversion over 12 weeks of treatment (n=30, 18 events; log-rank p=0.144; restricted mean survival time difference +1.1 weeks for the high-burden group). (B) Composite unfavourable treatment outcome (n=34, 7 events; log-rank p=0.400).

Sensitivity analyses tested whether results depended on influential participants or the cut-off. We excluded S0224_01 (a 50/50 lineage 2 mixture) and S0186_01 (lowest bacterial load, 312 genome copies, where LF calls may be less reliable), and replaced the AIC-optimal threshold of 25 variants, with the cohort median (>31). Results were similar (culture conversion log-rank p=0.162 and 0.139 for the exclusions, 0.147 for the median split; composite p=0.367 for each exclusion, 0.271 for the median split). Higher within-host genetic diversity measured directly from sputum showed a directional association with slower culture conversion and a higher proportion of unfavourable outcomes, but neither association was statistically significant. Larger cohorts are needed to evaluate the potential prognostic relevance of these findings.

## Discussion

In this prospective cohort of patients initiating treatment for rifampicin-susceptible pulmonary tuberculosis in South Africa, direct-sputum whole-genome sequencing (dsWGS) revealed substantial Mtb diversity that was not fully represented in matched culture-derived genomes. Mixed-lineage populations were detected in 63.7% of baseline sputum specimens compared with 6.2% of matched cultures, with reduced lineage diversity following culture observed in 58.4% of pairs. In contrast, genome-wide variant concordance remained high among pairs with concordant single-major-lineage classifications. These findings suggest that culture-derived sequencing can preserve much of the dominant genomic background while incompletely representing the bacterial population diversity detectable directly in sputum.

Previous studies have demonstrated that in vitro culture can alter the representation of Mtb populations, with greater genomic diversity detected directly in sputum and differences in mixed-infection recovery influenced by strain genotype and culture conditions (6–8). Our findings extend these observations through a systematic comparison of 113 paired baseline specimens from patients with rifampicin-susceptible pulmonary TB. The current study was conducted in Worcester, within the Cape Winelands district of South Africa, where historically high TB incidence has been documented (28). This substantial background burden may increase opportunities for repeated exposure to different Mtb strains, potentially contributing to the mixed-lineage diversity observed in our cohort. However, differences in sequencing approaches, sampling and the sensitivity of minority-lineage detection may also influence the observed frequency of mixed-lineage infections.

Previous paired sequencing studies have reported high sputum–culture variant concordance following careful bioinformatic filtering, with findings reproduced across several epidemiological settings (13). Similarly, 92.8% of pair-specific variant observations in our strain-controlled subset were shared following filtering and bidirectional recovery. However, our broader analysis revealed substantial differences in lineage composition, which were not apparent in the previously reported concordant pairs. Although previous studies have included geographically distinct populations, differences in cohort composition, mixed-infection prevalence, sequencing methods and culture conditions may influence the extent to which these findings are generalisable to settings with frequent mixed-lineage infection. Importantly, our strain-controlled analysis excluded pairs with differing major-lineage compositions, whereas the greatest differences in the broader cohort occurred at the lineage level. Thus, high genomic concordance within a shared major lineage does not necessarily imply equivalent representation of minority bacterial populations.

Differences in variant recovery were concentrated among variable variants, with additional low-frequency variation detected in sputum and higher culture allele frequencies among most shared variants that were variable in at least one specimen. Given the challenges associated with reliable low-frequency variant detection, our analysis incorporated benchmark-informed variant calling and an empirical error model to reduce false-positive SNP calls, alongside genomic masking and additional quality filtering (23, 29). These measures strengthen confidence in the observed differences, although residual discordance cannot be attributed exclusively to culture-associated selection, as sequencing depth, sampling variability and variant-detection thresholds may also contribute (13).

Our technical reproducibility analyses provide additional context for interpreting these findings. Repeated sequencing of the same DNA extract showed consistent dominant phylotype assignments, whereas low-frequency SNP concordance was lower. Independently collected same-day sputum specimens showed greater variation in lineage composition, suggesting that individual expectorates may sample different components of the bacterial population. Similarly, three longitudinal specimen pairs demonstrated relatively conserved fixed SNP profiles but greater differences in detectable low-frequency variation. These observations highlight the importance of distinguishing sequencing reproducibility from biological sampling variability when investigating within-host diversity. Previous studies have also demonstrated that Mtb populations can be spatially heterogeneous within the host and may not be fully represented by individual sputum specimens (30, 31).

Variants in drug-resistance-associated genes were largely conserved among the 38 strain-controlled pairs, with no sputum-unique WHO Grade 1 or 2 mutations identified. However, among mixed-lineage infections, Grade 1 or 2 mutations were detected in eight sputum specimens compared with three matched cultures, with additional sputum-only observations concentrated in pairs showing reduced lineage diversity following culture. These findings raise the possibility that differences in strain representation may influence the detection of resistance-associated variation. Nevertheless, the presence of catalogue-associated mutations does not independently establish phenotypic resistance, and our results do not demonstrate that these differences would have altered clinical management.

Exploratory clinical comparisons identified associations between mixed-lineage detection and alcohol-use measures, baseline culture time to positivity and radiographic lung involvement. These unadjusted findings require cautious interpretation. Among concordant single-major-lineage infections, higher baseline low-frequency (LF) variant burden showed directional associations with slower culture conversion and more unfavourable outcomes, but neither association was statistically significant. Analyses treating LF burden as a continuous variable also showed no evidence of association, while the threshold-based comparisons were limited by the small number of events and data-driven threshold selection. Previous work in the broader TRUST II cohort identified associations between within-host variant burden and treatment outcomes, although differences in specimen type and analytical approaches limit direct comparison, and the overlapping study populations preclude independent validation (14). Our findings therefore do not establish the prognostic value of direct-sputum LF burden, but suggest that further investigation in larger cohorts may be warranted.

### Limitations

Several considerations should be taken into account when interpreting these findings. Direct-sputum sequencing success was influenced by bacterial burden, with 139 of 218 clinical sequencing reactions meeting the predefined quality criteria. The resulting paired cohort was therefore enriched for specimens with sufficient bacterial DNA, which may affect the generalisability of our findings to specimens with lower bacterial burdens. Although technical replicates and process controls supported the reproducibility of dominant lineage assignments, minority-lineage detection was not independently confirmed for every specimen. Furthermore, the strain-controlled analysis was restricted to 38 pairs and could not exclude mixtures of distinct strains belonging to the same major lineage. Despite extensive quality filtering, differences in specimen processing, sequencing depth and variant-detection thresholds may contribute to sputum–culture discordance alongside biological effects of culture. Only one routine liquid-culture approach was evaluated, and the lineage permutation analysis did not incorporate initial strain proportions. Finally, limited longitudinal sampling, multiple exploratory clinical comparisons and the small number of treatment-outcome events restrict conclusions regarding within-host evolution and the clinical relevance of the observed diversity.

Overall, our findings highlight the complementary value of direct-sputum and culture-derived WGS. While culture remains important for generating high-quality genomes and supporting phenotypic investigations, its ability to represent the bacterial diversity present in sputum may vary across epidemiological settings and study populations, particularly where mixed-lineage infections are frequently detected. Considered alongside previous reports of high sputum–culture concordance, our findings emphasise the importance of accounting for within-sample population diversity when interpreting culture-derived genomic data.

### Conclusion

In conclusion, direct-sputum WGS revealed substantial Mtb lineage and low-frequency variant diversity that was not fully represented by matched culture-derived genomes in this high-burden South African cohort. Despite high overall genomic concordance, differences in detectable bacterial populations highlight that culture-derived sequencing may incompletely represent within-sample diversity, with the extent of these differences potentially varying across epidemiological settings. These findings support the complementary use of direct-sputum and culture-derived WGS for investigating within-host Mtb diversity. Future research should focus on optimising specimen processing and genomic recovery, improving the accuracy and reproducibility of minority-variant detection, and determining whether diversity measured directly from sputum provides clinically relevant information on treatment response. Together, these efforts will help establish the biological significance and potential clinical utility of genomic diversity that is not fully captured by conventional culture-based sequencing.

## Supporting information

Datafile - S1

