## Supplementary methods and figures for "Direct Sputum Sequencing Reveals *Mycobacterium tuberculosis* Diversity Underrepresented by Culture in a High TB-Burden Setting"

#### Specimen processing and DNA extraction

Clinical sputum specimens were decontaminated using the BD BBL™ MycoPrep™ kit according to the manufacturer’s instructions. Decontaminated sediments were divided for multiple purposes: a portion was used for Acid-Fast Bacilli (AFB) Ziehl–Neelsen (ZN) staining to assess bacillary load, and another portion was inoculated into MGIT tubes to establish primary liquid cultures. Residual sediment was retained for DNA extraction and subsequent direct sputum whole-genome sequencing (dsWGS) using the TWIST capture protocol as outlined previously (1). Primary MGIT cultures flagged as positive were screened for contamination through repeat ZN staining and inoculation onto blood agar. Confirmed MGIT cultures were subcultured in 7H9 liquid medium to generate sufficient biomass for DNA extraction and culture-based WGS. In parallel, subcultures were used for first-line phenotypic drug susceptibility testing (pDST) performed on the BACTEC MGIT system according to manufacturers instructions (2, 3). To monitor culture growth, DNA extraction, and sequencing performance, two reference *Mtb* H37Rv cultures were inoculated into 7H9 medium and processed in parallel with clinical isolates through all subsequent culture, inactivation, DNA extraction, and sequencing steps.

All procedures involving live *Mtb* were performed within a biosafety level 3 (BSL-3) laboratory in accordance with institutional safety protocols, and both direct sputum and culture samples were heat-inactivated at 80 °C for 60 minutes prior to DNA extraction (1, 4). DNA extraction from direct sputum samples was performed using a modified column‑based approach with the Zymo Clean & Concentrator‑25 kit, as previously described (1). For culture‑derived DNA, extraction was carried out either using the same Zymo‑based protocol or by a standard CTAB chemical lysis method adapted from established *Mtb* DNA extraction workflows (1, 4).

#### Quantitative PCR

Quantitative PCR (qPCR) was used to estimate *Mtb* burden in each decontaminated sputum sediment prior to target capture and enrichment to guide pooling during hybrid capture as previously described. The assay targeted the single-copy *Mtb*–specific gene *Rv2341*, using the primer pairs Rv2341-F (5′-GCCGCTCATGCTCCTTGGAT-3′) and Rv2341-R (5′-AGGTCGGTTCGCTGGTCTTG-3′). Amplification and fluorescence detection were performed in the SYBR/FAM channel on a CFX96 Opus Real-Time PCR Detection System (Bio-Rad, USA). For quantification, standard curves were generated using five concentrations of previously extracted H37Rv DNA, prepared by serial dilution to yield input amounts of 10, 1, 0.1, 0.01, and 0.001 ng per reaction. The mean cycle threshold (Ct) value for each sample was recorded and, together with the standard curve, used to estimate the approximate *Mtb* genome copy number per sample (1, 5).

#### Library preparation and target capture and enrichment

A set of custom-designed and previously validated target capture and enrichment DNA bait probes (Twist Bioscience, USA) was used for the enrichment and sequencing of direct patient samples. Library preparation for all samples was performed using the Twist enzymatic fragmentation and universal adapter system, following the manufacturer’s protocol with two modifications. Given the limited DNA input concentrations, and in accordance with Twist’s recommendations, the adapter concentration was halved, and the number of amplification cycles using Twist UDI primers was increased from 8 to 10 cycles (1). Two versions of the previously validated custom Twist capture panel were used during the study. The original panel was subsequently modified to reduce probe representation targeting the highly conserved *rrs* and *rrl* genes, which had shown disproportionate read enrichment and potential off-target hybridisation. This modification was intended to improve sequencing efficiency and coverage across the remaining targeted genomic regions. The distribution of samples sequenced using each panel version is provided in Supplementary Data – S1.

Libraries were pooled in sets of eight, grouped based on similar genome copy numbers as determined by prior qPCR quantification, to minimize capture bias associated with pooling high-load and low-load samples. Final pooled libraries were sequenced using either an Illumina MiniSeq with the High Output Reagent Kit (300 cycles) or an Illumina NextSeq 1000/2000 with the P2 Reagent Kit (300 cycles) at the Stellenbosch University Centre for Epidemic Response and Innovation (CERI). Culture samples were shipped for sequencing at the Leibniz Lung Research Center (Borstel, Germany) according to the centres standard procedures.

#### Bioinformatics – Core analysis

Initial quality assessment of raw sequencing data was performed using FastQC (v0.11.9) (6). Adapter trimming, quality filtering, and per-read quality pruning were conducted using fastp (v0.23.4) (7). Following quality control, taxonomic classification was performed using Kraken2 v2.1.3 to identify reads originating from *Mtb* (NCBI taxID 1773 (8)). Reads classified as *Mtb* were subsequently extracted using KrakenTools to reduce downstream interference from contaminating background organisms (8, 9). *Mtb* classified reads were aligned to the H37Rv reference genome (NC_000962.3 ) using bwa-mem2 (v2.2.1) with an increased seed length parameter (-k 80) to improve mapping stringency in the presence of residual non-target DNA (10). Resulting SAM files were converted to BAM format using samtools (v1.20) view (-b -h) while retaining primary alignments and removing secondary and supplementary alignments known to cause spurious variant calls in direct from sputum data (-F 256 -F 2048)  (11, 12). Read group information was added using samtools addreplacerg (12). PCR duplicates were subsequently identified and removed using Picard Tools MarkDuplicates (REMOVE_DUPLICATES=true) (v3.1.1) to minimise amplification bias (13). Per-base coverage depth was computed using samtools depth, and GC bias metrics were collected using Picard CollectGcBiasMetrics for quality assessment (12, 13).

Following deduplication, variant calling was performed using FreeBayes (v1.3.6) (14) with parameters optimised for haploid bacterial genomes (--ploidy 1, --min-mapping-quality 30, --min-base-quality 30, --min-alternate-count 2, --min-alternate-fraction 0.01, --report-all-haplotype-alleles). Raw variant calls were then decomposed into substitutions and indels: indels and complex variants were realigned using vcfwave (vcflib v1.0.9), while SNPs and multi-nucleotide polymorphisms (MNPs) were extracted using bcftools view (v1.20) (12, 14, 15). The resulting calls were recombined and sorted. A custom script was applied to correct haplotype-level allelic support annotations introduced during all-haplotype reporting. Functional annotation was performed using SnpEff (v5.2) with the Mycobacterium_tuberculosis_h37rv database prior to multiallelic splitting to ensure that multi-nucleotide variants (MNVs) were annotated prior to splitting (16). Multiallelic records were then normalised and split using bcftools norm (v1.20), and variants were filtered using bedtools subtract (v2.31.0) (17) with a custom BED file (Supplementary File - exclusion regions) excluding 471,463 bp (10.7%) of the 4,411,532 bp H37Rv genome comprising PE/PPE genes, rRNA loci, and regions of high sequence homology, poorly mappable loci, and genomic intervals prone to cross-species misalignment. Following splitting, per-allele observation counts (AO) were corrected to reflect the decomposed records. Variant fields including position, alleles, quality, functional annotations (gene, HGVS coding, and HGVS protein changes), and all INFO-level attributes were extracted into tabular format using SnpSift  (v5.2) extractFields (16).

#### Variant quality filtering

For SNP filtering, per-variant alignment statistics were calculated from deduplicated BAM files, including local sequencing depth, coverage relative to a rolling neighbourhood average, soft-clipped base counts, alternate allele observations and discordant read counts. Variants were required to meet a minimum mapping quality mean (MQM) of 40, at least five alternate allele observations (AO ≥5), and bidirectional strand support (SAF ≥2 and SAR ≥2). A minimum allele frequency of 0.05 was required, with additional depth-dependent AF thresholds applied as summarised in Supplementary Table – S1. Variable SNPs were subsequently evaluated using the previously described empirical error model (18, 19). For indel filtering, CIGAR strings from reads overlapping candidate indel positions were examined to assess alignment support. Adjusted allele frequencies were calculated using only reads with alignment start and end positions at least 10 bp from the indel boundaries, excluding reads containing soft-clipped bases. A Mann–Whitney U test comparing alignment start and end positions of indel-supporting and reference-supporting reads was used to identify potential fixed indels misrepresented as heterogeneous. Duplicate indel records were removed, and the remaining indels were subjected to the depth-dependent AF thresholds specified in Supplementary Table - S1.

#### Lineage classification and concordance analysis

Lineage assignments were generated using TBTypeR (v25.01.0; ploidy 1). Variants at curated phylogenetic marker positions were called using bcftools mpileup (-q 1, -Q 10, -d 200) and bcftools call (--ploidy 1, -A, -m, --prior 1e-2, -C alleles). Marker positions overlapping the predefined low-confidence genomic mask were excluded before lineage assignment (20). For paired comparisons, lineage classifications were represented as sets of detected major lineages. Concordant single-lineage pairs contained the same single major lineage in both specimens, whereas concordant mixed-lineage pairs contained identical sets of two or more major lineages. Reduced diversity in culture was assigned when the culture lineage set was a proper subset of the direct-sputum set, and reduced diversity in sputum when the direct-sputum set was a proper subset of the culture set. Pairs with differing lineage sets that did not meet either subset relationship were classified as discordant. To evaluate lineage-neutral dropout, the 208 major-lineage observations detected across the 113 baseline direct-sputum specimens formed the sampling pool. For each of 200,000 simulations, 120 lineage observations, corresponding to the total observed in matched cultures, were sampled without replacement. Simulated counts for Lineages 2, 3 and 4 were compared with their observed culture counts to evaluate departures from lineage-neutral sampling.

#### Variant concordance and recovery

For each of the 38 eligible baseline sputum–culture pairs, final quality-filtered SNP and indel call sets were compared using exact genomic position, reference allele and alternate allele combinations. Fixed and variable variant calls were combined for concordance assessment while retaining their original classification. Variants detected in both specimens were classified as shared, whereas variants detected in only one specimen were provisionally classified as direct sputum-unique or culture-unique.

Initially unique variants underwent bidirectional recovery by interrogating pre-filter sequencing evidence in the matched specimen. The documented recovery criteria required a minimum raw depth of two reads, at least one alternate-supporting read and a raw allele frequency ≥0.01. Recovered variants were further categorised as high confidence (raw depth ≥10 and raw AF ≥0.10) or medium confidence. Where qualifying evidence was identified, the corresponding variant was added to the matched specimen's rescued call set, with allele frequency and depth obtained from the recovered evidence.

Final concordance classifications were determined from the rescued call sets. Variants present in both specimens after recovery were classified as shared, including variants that had not independently passed all initial quality filters in both specimens. Variants detected in only one rescued call set remained specimen-unique. These classifications represented differences in variant detection rather than independent confirmation of the reference genotype in the opposing specimen. Variant counts represented pair-specific observations, such that the same genomic variant could contribute separately in multiple specimen pairs. Allele-frequency differences among shared variants were calculated for corresponding alleles in each specimen and summarised at the specimen-pair level for statistical comparison.

#### Genomic distribution and annotation-level analysis of specimen-unique variants

Following variant filtering and bidirectional raw-read recovery, the genomic positions of direct sputum-unique and culture-unique variants were extracted from the final concordance dataset. Variant observations were assigned to their corresponding genes or genomic regions using the annotations generated during variant processing. Genome-wide distributions were visualised using 500 position-based bins across the H37Rv reference genome. For each specimen type, annotations were ranked according to the number of specimen-unique variant observations, and the 15 most frequently represented annotations were selected for visualisation. Counts represented pair-specific variant observations rather than distinct genomic mutations, such that an identical variant detected across multiple specimen pairs contributed an observation for each pair.

#### Drug-resistance-associated variant analysis

##### Strain-controlled baseline pairs

Drug-resistance-associated variants were extracted from the final rescued variant-call sets for the 38 concordant single-lineage baseline sputum–culture pairs. Variants within the curated resistance-associated gene set were classified as shared, direct sputum-unique or culture-unique according to their detection in matched specimens. Concordance was summarised using pair-specific variant observations, while distinct genomic mutations were identified by exact genomic position, reference allele and alternate allele. Drug-resistance associations were assessed through exact mutation-level matching against the WHO Catalogue of mutations in *Mycobacterium tuberculosis* complex and their association with drug resistance (third edition; 2026). Grade 1 and Grade 2 classifications were considered evidence of an association with resistance, Grade 3 indicated uncertain significance, and Grades 4 and 5 indicated mutations classified as not associated with resistance under the catalogue framework (21).

##### Mixed-lineage baseline pairs

An exploratory resistance-associated variant screen was conducted in 72 baseline sputum–culture pairs classified as mixed-lineage by direct sputum sequencing. Final quality-filtered fixed SNPs, variable SNPs retained following error-model filtering (classification probability ≥0.67), and quality-filtered fixed and variable indels were screened independently in direct sputum and culture-derived sequencing datasets. Variants were matched against the WHO 2026 catalogue using exact genomic position, reference allele and alternate allele, retaining Grade 1 and Grade 2 resistance-associated classifications. Where a genomic variant had multiple drug-specific catalogue classifications, it was counted as a single variant observation within each specimen. Detection concordance and allele frequencies were compared between matched specimen types. Reported variants were subsequently checked against their original sequencing outputs and corresponding WHO catalogue entries.

#### Technical reproducibility, sputum sampling variability, and process controls

##### Technical replicate reproducibility

Technical replicate analyses were restricted to specimens with at least two independent sequencing runs from the same DNA extract meeting the predefined sequencing quality criteria. The final analysis comprised seven specimen sets, including four with three qualifying sequencing runs and three with two qualifying runs. All possible pairwise comparisons were performed within each specimen set, including all three combinations for specimens with three sequencing runs, producing 15 pairwise comparisons from 18 sequencing runs. Variant reproducibility was assessed using final quality-filtered call sets. Fixed SNPs were obtained from the filtered variant outputs, variable SNPs from the error-model-filtered outputs, and fixed and variable indels from their corresponding quality-filtered outputs. Variant identities were compared using genomic position, reference allele and alternate allele. Jaccard similarity was calculated as the number of variants shared between two sequencing runs divided by the total number of distinct variants detected across the two runs. Similarity was evaluated for all SNPs and separately for fixed and variable SNPs, with additional comparisons of indel and combined variant sets. Phylotype reproducibility was assessed by comparing TBTypeR major-lineage and phylotype assignments between sequencing runs. Concordance was evaluated using the presence or absence of detected lineages and phylotypes, independently of differences in their estimated proportions.

##### Same-day sputum sampling variability

Independent sputum specimens collected during the same study visit were evaluated separately from technical sequencing replicates. Six A/B specimen pairs in which both samples passed sequencing quality control were included. Major-lineage classifications and complete TBTypeR phylotype sets were compared between paired specimens using Jaccard similarity.

Phylotype concordance was assessed using exact TBTypeR labels and a hierarchy-aware comparison in which related parent and descendant phylotype assignments were harmonised to represent the same phylogenetic branch. This allowed differences in sublineage resolution to be distinguished from detection of unrelated phylotypes. Changes in the presence and relative representation of major lineages and minority phylotypes were assessed descriptively.

##### H37Rv process controls

Two H37Rv reference process controls were evaluated using their sequencing quality metrics, filtered variant profiles and TBTypeR lineage assignments. Concordance between controls was examined to assess technical consistency and identify unexpected genomic or phylogenetic signals potentially indicative of contamination or systematic processing artefacts.

#### Exploratory clinical correlates of mixed-lineage infection

The analysis was restricted to the 113 participants in the corrected baseline paired-sequencing cohort, comprising 72 mixed-lineage and 41 single-lineage infections according to direct-sputum WGS. Each participant contributed one baseline lineage classification. Clinical information was linked by participant identifier to the TRUST II clinical extract dated 1 July 2026.

The descriptive comparisons followed the variable framework used in the original clinical analysis. Characteristics included sex, education, incarceration history, body mass index category, race/ethnicity, employment status, smoked substance use, age category, household hunger, depression risk, HIV status and HIV-related treatment history, alcohol-use measures, previous tuberculosis, baseline sputum microscopy, liquid-culture time to positivity and chest radiographic findings.

Alcohol-related measures included heavy alcohol use, problem alcohol use, Alcohol Use Disorders Identification Test (AUDIT) category and baseline phosphatidylethanol (PEth) concentrations categorised as <50, 50–200 or >200 ng/mL. Baseline smear positivity was additionally assessed using the combined TB programme or TRUST microscopy classification. Time to positivity was obtained from the whole-day measurement for the first sputum specimen. Percentage lung involvement was derived from the first chest radiograph.

Categorical variables were summarised as frequencies and percentages and compared using Pearson's chi-square tests without Yates' continuity correction. Continuous variables were summarised as medians and interquartile ranges and compared using two-sided Mann–Whitney U tests. Missing values were excluded separately for each comparison, and available sample sizes were reported. No imputation, multivariable adjustment or multiple-testing correction was performed. All analyses were exploratory.

### Supplementary figures

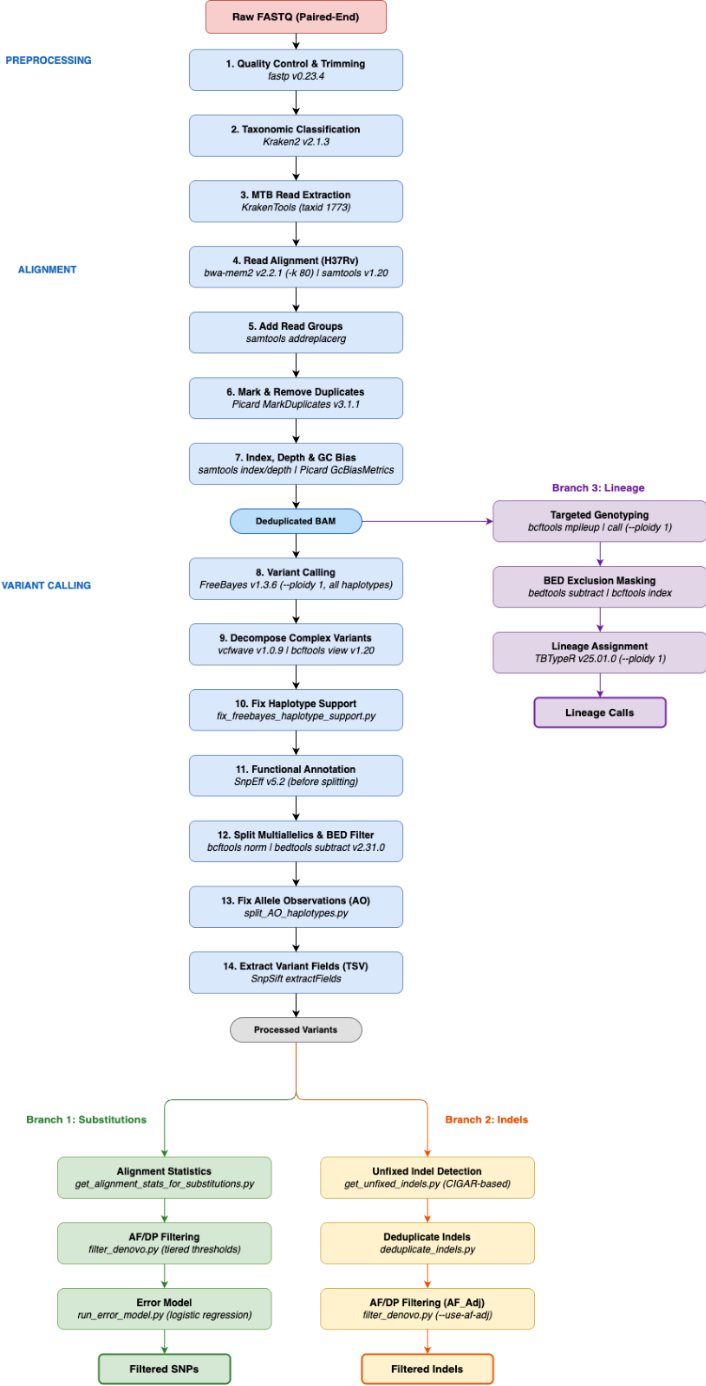

**Supplementary Figure - S1: Bioinformatics workflow for whole-genome sequencing analysis.** Overview of sequencing read processing, quality control, alignment to the H37Rv reference genome, variant calling and filtering. Downstream analyses followed separate branches for SNP filtering, indel filtering and phylogenetic lineage classification using TBTypeR.

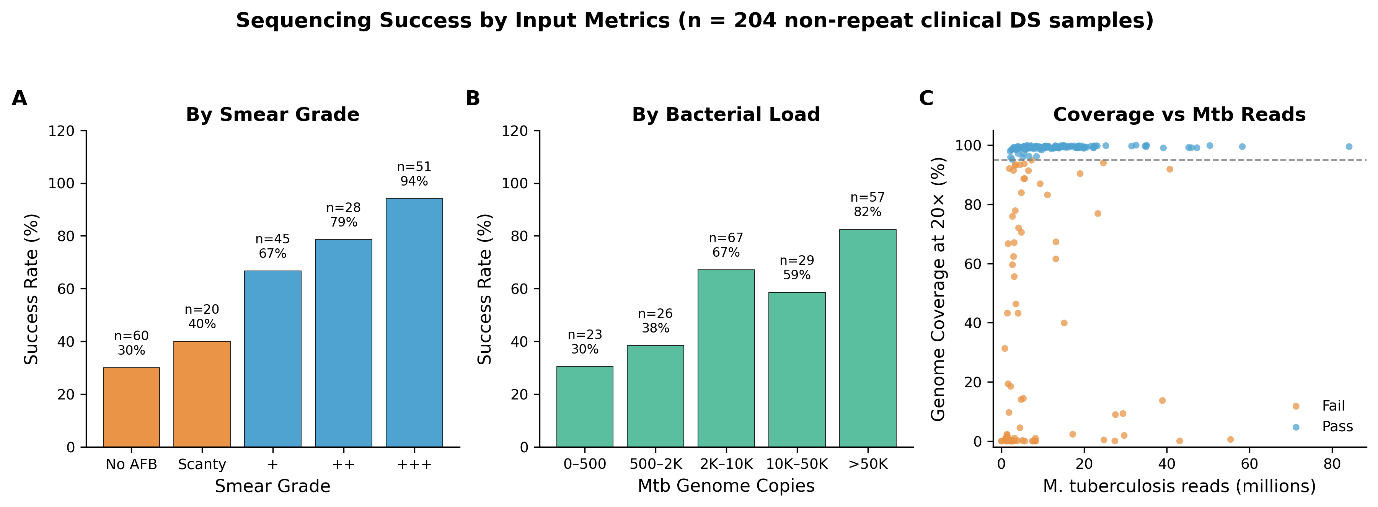

**Supplementary Figure – S2: Sequencing success by bacterial burden.** Sequencing success stratified by (A) smear microscopy grade and (B) Mtb genome copy number. (C) Relationship between Mtb-classified read yield and genome coverage, with samples classified as pass or fail according to the predefined threshold of ≥95% genome coverage at ≥20× depth. Panel B excludes two samples without quantitative Mtb-copy estimates.

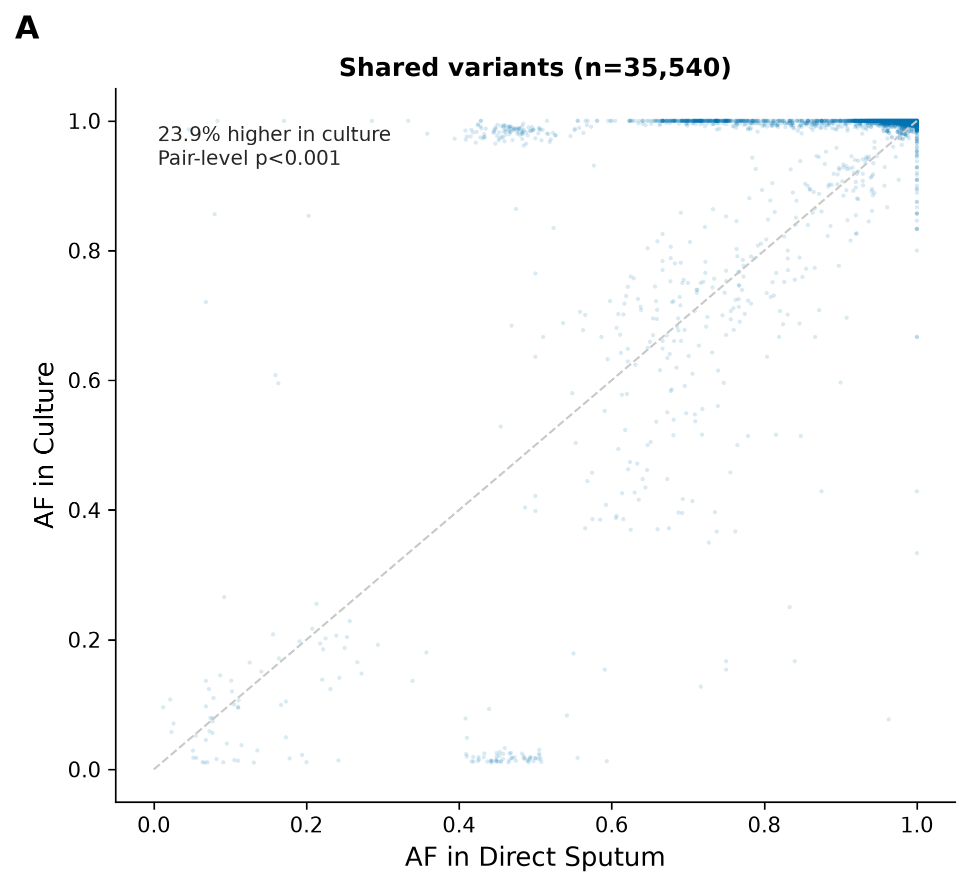

**Supplementary Figure – S3: Allele-frequency concordance among variants shared between direct sputum and culture-derived WGS.** Scatterplot of paired allele frequencies among 35,540 shared variant observations from 38 concordant single-lineage baseline pairs. The dashed diagonal indicates equal allele frequency in both specimen types. Points above the diagonal indicate higher allele frequencies in culture, whereas points below indicate higher frequencies in direct sputum. Statistical significance was assessed using a paired, specimen-level Wilcoxon signed-rank test

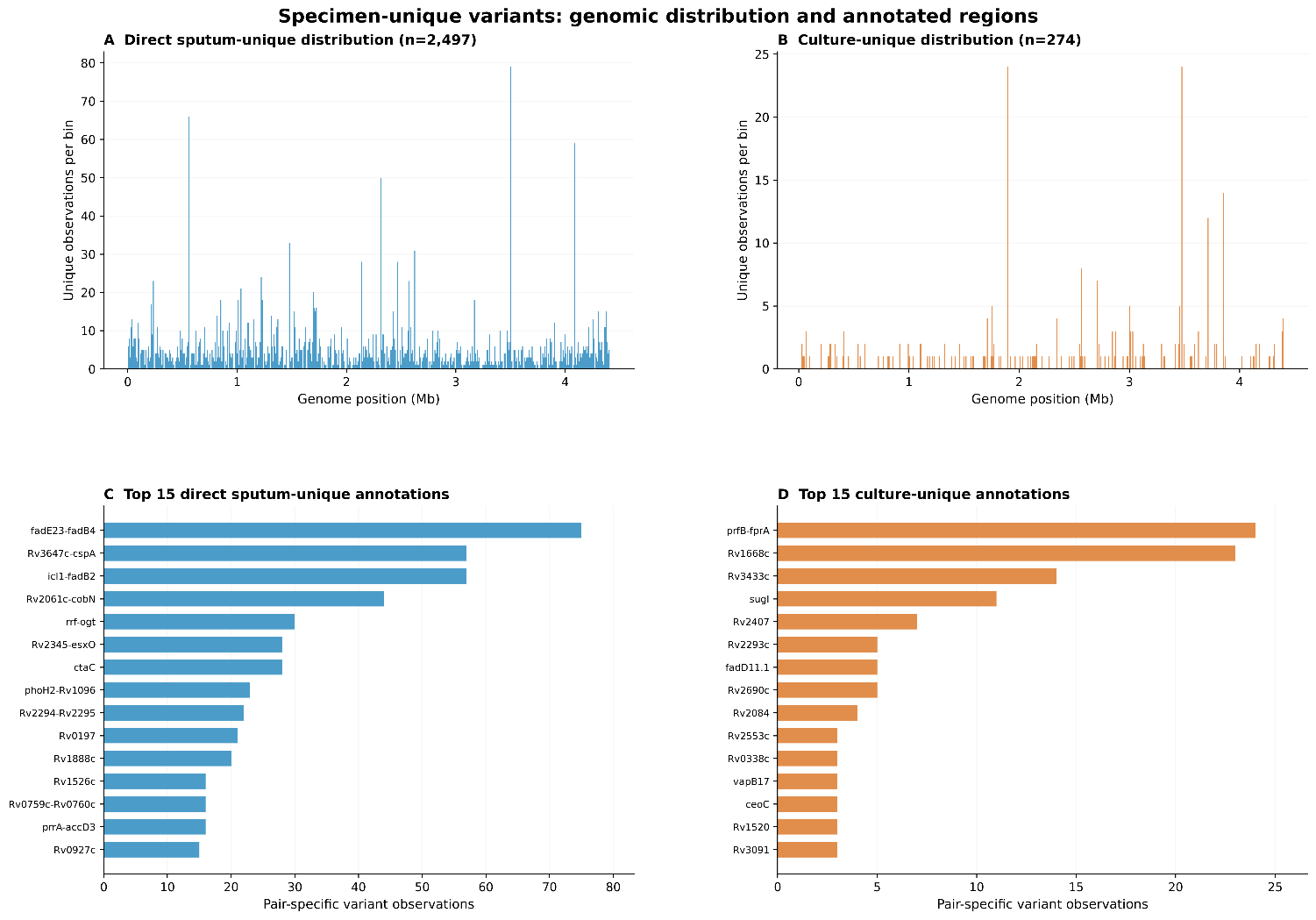

**Supplementary Figure - S4: Genomic distribution and annotation-level patterns of specimen-unique variants in 38 baseline sputum–culture pairs.** (A–B) Genome-wide distribution of direct sputum-unique (n = 2,497) and culture-unique (n = 274) variant observations, respectively. (C–D) The 15 gene or genomic-region annotations with the highest numbers of direct sputum-unique and culture-unique variant observations, respectively. Counts represent pair-specific variant observations, meaning that the same genomic variant may contribute observations from multiple specimen pairs.
